# Longevity Relatives Count score identifies heritable longevity carriers and suggests case improvement in genetic studies

**DOI:** 10.1101/609891

**Authors:** Niels van den Berg, Mar Rodríguez-Girondo, Kees Mandemakers, Angelique A.P.O. Janssens, Marian Beekman, P. Eline Slagboom

## Abstract

Longevity loci represent key mechanisms of a life-long decreased mortality and decreased/compressed morbidity. However, identifying such loci is challenging. One of the most plausible reasons is the uncertainty in defining long-lived cases with the heritable longevity trait amongst long-living phenocopies. To avoid phenocopies, family selection scores have been constructed but these have not yet been adopted as state of the art in longevity research. Here we aim to identify individuals with the heritable longevity trait by using current insights and a novel family score based on these insights. We use a unique dataset connecting living study participants to their deceased ancestors covering 37,825 persons from 1,326 five-generational families, living between 1788 and 2019. Our main finding suggests that longevity is transmitted for at least 2 subsequent generations only when at least 20% of all relatives are long-lived. This proves the importance of family data to avoid phenocopies in genetic studies.

## Main

In contrast to the low heritability of human lifespan^1–4^, human longevity is strongly heritable as illustrated by the familial clustering of survival into extreme ages^5–17^. Identifying longevity loci is important because these loci likely represent key mechanisms of a life-long decreased Mortality^15, 16^, decreased morbidity^9, 12, 18^ and compression of morbidity towards the end of the lifespan^19–21^. Currently, genome wide linkage and association studies (GWAS) identified a limited number of loci promoting longevity^22–31^, for example the *APOE* and *F0X03A* genes (more details can be found in current review papers^22, 23, 30^). However, many of the identified loci could not be replicated in independent studies as yet. In addition, the largest and most recent longevity GWAS, based on cases belonging to the top 10% oldest survivors, again only replicated association of the APOE locus^32^.

One of the main reasons for the limited success of longevity genetic studies^24–26, 31–34^ the uncertainty in defining the heritable longevity trait itself^1, 16^. Given the increased life expectancy of the past 200 years due to non-genetic factors (improved hygiene, nutrition and medication) there are likely many phenocopies among the long-lived cases selected for our genetic studies^35, 36^. The presence of phenocopies is illustrated by the increase of centenarians in the United States between 1994 and 2012 from 1 in 10,000 to 1 in 5,000^37^. To avoid phenocopies, family selection scores, such as the Family Longevity Selection Score (FLoSS) and the Family Excess Longevity (FEL) score have been constructed^38, 39^. The use of such scores is substantiated by novel studies which showed that that including family history information can provide valuable information about an individual’s genetic liability for a trait and is likely to increase the power to detect genetic^40–42^. The scores focus, in different ways, on selecting multiple family members with the same trait^15, 38, 39, 43, 44^ and usually focus on a single group of relatives, such as parents^15, 43^ or siblings^39^ of cases.

As the definition of heritable longevity was not yet established, the construction and application of the family selection scores have not yet been adopted as state of the art in longevity research. As such, the majority of genealogical^5, 6, 10–14, 45^ and genetic studies^24–26, 31–34^ focus only on single, and thus including sporadic, long-lived individuals (singletons), with some exceptions focusing for example on parental age^28, 29^ or multiple siblings^7, 25^ . In previous work, we showed that longevity defined as top 10% survivors or more extreme is transmitted to subsequent generations^16^. With this, a consistent definition of longevity was provided that is also adopted in the largest longevity GWAS up to now^32^. In addition, we showed that every additional long-lived relative independently contributes to the survival advantage of study participants, according to their genetic distance^16^. As such, there is room to incorporate these novel insights into family selection scores to gain knowledge about the extent that longevity needs to cluster in families in order to include individuals with the heritable longevity trait and increase the power of genetic studies.

Here, we aim to establish the proportion of ancestral blood relatives that should be long-lived (top 10% survivors of their birth cohort or more extreme) in order to observe a survival advantage in their descendants and incorporate these insights into a novel family score to define cases with the heritable longevity trait for inclusion in genetic studies. For our analyses we use the data available in the Historical Sample of the Netherlands (HSN) for the period between 1860 and 1875 which is based on Dutch citizens^46–48^. We primarily identify cases who died beyond 80 years (N=884, on average top 10% survivors of their birth cohort), allowing us to select on more extreme ages at death, and controls who died between 40 and 59 years (N=442). We extend this filial (F) 1 generation data with a parental and 3 descendant generations of individual life course and mortality data and refer to the data as the HSN case/control dataset. We subsequently exclude groups with high rates of missing mortality information and where the majority was still alive (Supplementary Figure 4). This study covers 37,825 persons from 1,326 three-generational families (F1-F3) and contains FI index persons (IPs), 2 consecutive generations of descendants (F2-F3) and 2 generations of spouses (F2-F3) (Table 1). The dataset is unique in that it covers multiple generations and connects alive persons to at least two generations of deceased ancestors.

**Table 1:** Overview study sample for groups in all generations based on the proband and F3 perspective

| Role | Number | Deceased (%) | Alive (%) | Female (%) | Range Birth cohort | Mean age (sd) | Median age (sd) | missing_age (%) |
| --- | --- | --- | --- | --- | --- | --- | --- | --- |
| Cases (Original design) |  |  |  |  |  |  |  |  |
| F1 IPs | 884 | 884 (100) | 0 (0) | 422 (50) | 1860-1875 | 85.79 (4.59) | 84.99 (4.95) | 0 (0) |
| F2 descendants | 4916 | 4405 (90) | 11 (1) | 2435 (50) | 1879-1941 | 63.04 (31.11) | 75.51 (17.72) | 500 (9) |
| F2 spouses | 3899 | 1500 (38) | 16 (1) | 1504 (38) | 1873-1934 | 76.2 (15.09) | 78.78 (12.83) | 2383 (61) |
| F3 descendants | 9910 | 4869 (49) | 4146 (42) | 4733 (48) | 1901-1973 | 70.35 (19.54) | 74.77 (11.38) | 895 (9) |
| F3 spouses | 3431 | 1289 (38) | 792 (23) | 1963 (57) | 1900-1959 | 77.14 (11.31) | 79.25 (10.1) | 1350 (39) |
| F4 descendants* | 9001 | 746 (8) | 7172 (80) | 3937 (44) | 1922-1995 | 57.7 (10.68) | 58.21 (9) | 1083 (12) |
| Controls (Original design) |  |  |  |  |  |  |  |  |
| F1 IPs | 442 | 442 (100) | 0 (0) | 214 (48) | 1860-1875 | 51.71 (5.71) | 52.88 (6.21) | 0 (0) |
| F2 descendants | 2488 | 2202 (89) | 1 (<1) | 1217 (49) | 1881-1925 | 58.17 (32.49) | 71.72 (21.37) | 285 (11) |
| F2 spouses | 1877 | 690 (37) | 7 (<1) | 734 (39) | 1875-1935 | 76.02 (14.77) | 78.34 (13.76) | 1180 (63) |
| F3 descendants | 4761 | 2540 (53) | 1813 (38) | 2265 (48) | 1904-1966 | 69.39 (20.38) | 74.49 (11.36) | 408 (9) |
| F3 spouses | 1778 | 721 (41) | 376 (21) | 972 (55) | 1893-1965 | 76.54 (11.5) | 78.66 (10.47) | 681 (38) |
| F4 descendants* | 4710 | 387 (8) | 3744 (80) | 2099 (45) | 1871-1992 | 57.72 (11.17) | 58.37 (9.35) | 579 (12) |
| F3 perspective (Combined design) |  |  |  |  |  |  |  |  |
| F3 descendants | 14671 | 7409 (51) | 5959 (41) | 6998 (48) | 1901-1973 | 70.03 (19.82) | 74.68 (11.38) | 1303 (8) |
| F3 spouses | 5209 | 2010 (38) | 1168 (22) | 2935 (55) | 1893-1965 | 76.93 (11.38) | 79.07 (10.24) | 2031 (40) |
| F2 parents | 9728 | 6139 (63) | 23 (1) | 4137 (43) | 1873-1935 | 76.8 (13.4) | 78.9 (12.31) | 3566 (36) |
| F2 aunts & uncles | 7036 | 6382 (91) | 10 (1) | 3456 (49) | 1879-1941 | 61.81 (31.47) | 74.4 (18.67) | 644 (8) |
| F1 grandparents | 1181 | 1181 (100) | 0 (0) | 560 (47) | 1860-1875 | 74.88 (16.6) | 81.94 (9.72) | 0 (0) |

The Cases and Controls rows provide an overview of the groups of persons from the original case/control perspective of the data, described as part a. The F3 perspective rows provide an overview of the groups of persons from the perspective of F3 descendants, described as part b. mean and missing age refer to an unknown age at death or an unknown age at last observation. For the F0 and F1 groups we assume everyone is dead because the birth cohorts date back further than 120 years. From the F2 generations we requested Personal Records Data indicating if a person was still alive or not and if not, what the date of death was. The F1 IPs are the focal persons in the pedigrees as they are selected to be 80 years or older (cases) or to have died between 40 and 59 years (controls). \* indicates that the group is excluded for this study, sd refers to standard deviation.

## Results

### Outline

We analyzed the data across multiple steps (Supplementary Figure 5) in two phases. In the first phase, we used Standardized Mortality Ratios (SMRs) to compare the transmission of longevity for cases (died beyond 80 years) and controls (died between 40 and 59 years) as defined in the original approach (Figure 1A), focusing on the F1 index persons (IPs) and two generations of descendants.

**Figure 1:**
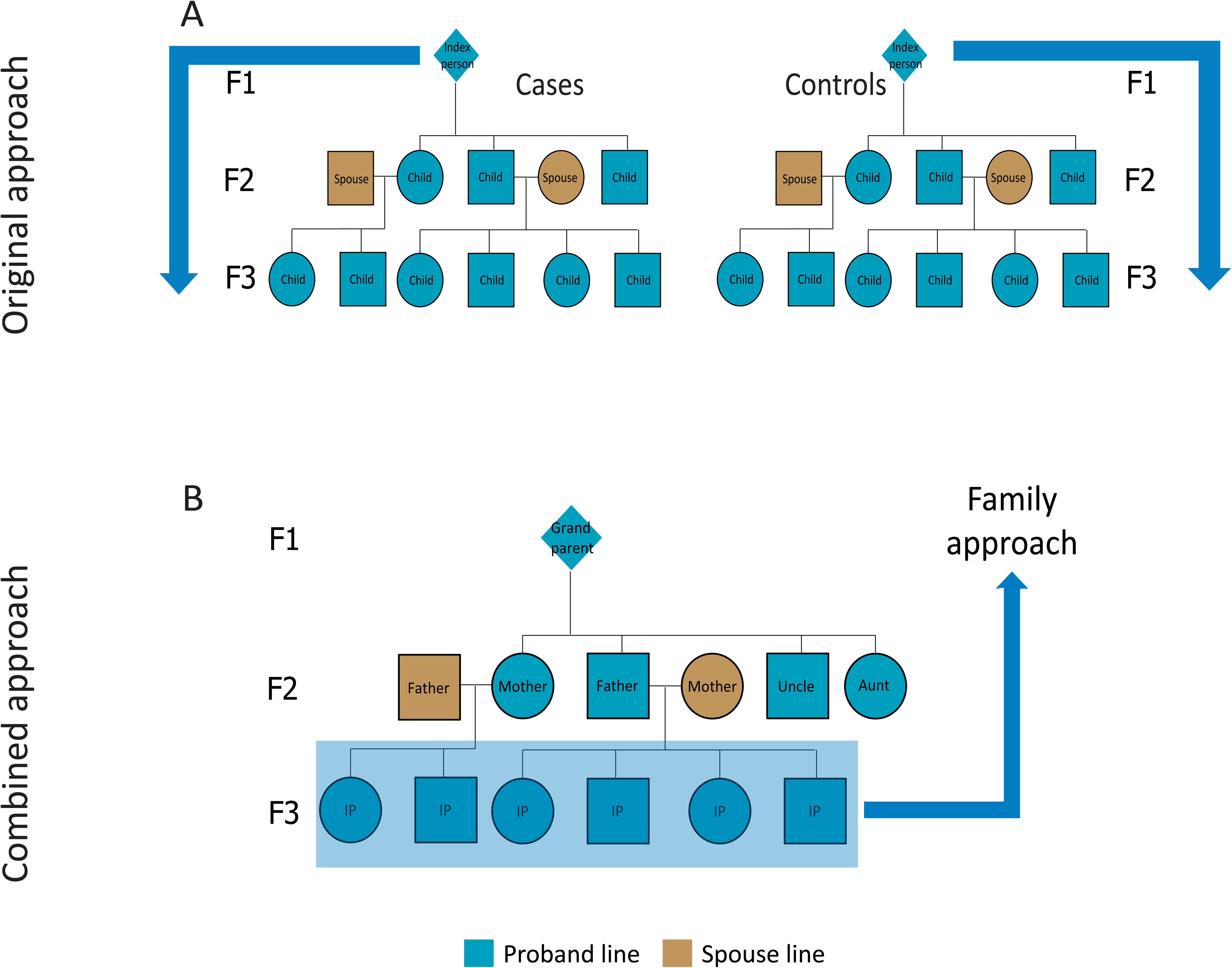
Pedigree overview of the data structure. This figure illustrates the two approaches; 1. the original approach and 2. the combined approach. The original approach refers to the case and control group based on the FI IPs where cases died at 80 years or older and controls died between 40 and 59 years (panel A). Panel B shows a pedigree of the data from the perspective of F3 children (combined approach). The combined approach refers to the dataset where we combined the cases and controls form the original design and constructed a new case and control group in the F3 descendants. To this end, F3 descendants with >30% long-lived ancestors were labeled as family cases and those without long-lived ancestors as family controls. F3 spouses were left out of this figure but this group was used to confirm a genetic enrichment in the F3 descendants.

In the second phase of our study (the combined approach), we combined original cases and controls and their descendants into one combined group and focused on the survival of the F3 descendants in relation to their F2 and F1 ancestral family members (Figure 1B). First, we constructed the Longevity Relatives Count (LRC) score. We used the LRC score to investigate the proportion of long-lived (top 10% survivors of their birth cohort) F1 and F2 ancestors required for F3 descendants to express a survival advantage compared to members of the same birth cohort and sex (family method, Figure 1B). On the basis of these observations we defined a new case and control group in F3, where we labeled F3 descendants with >30% long-lived ancestors as family cases and those without long-lived ancestors as family controls. Subsequently, these F3 family cases and controls were compared for their survival, that of their spouses (to investigate environmental influences), and for survival differences with the F3 descendants, selected to have at least one (singleton) long-lived ancestor or at least one average-lived ancestor. This means that they could have more than 1 long or average lived ancestor but we actively selected for the presence of only 1 such ancestor. Supplementary Figure 3A provides a conceptual overview of this selection. To this end, we selected either F3 descendants with at least one top 10% grandparent, at least one top 10% parent, or with grandparents who died between 40 and 59 years (their children (parents) resembled the general population). In a final step, we focused on the F3 descendants with at least one long-lived parent and calculated LRC scores within this F3 group to determine if parents transmitted their longevity more frequently if they were part of a long-lived (LRC≥0.30) family (Figure 1B). The analysis steps are summarized in Supplementary Figure 5 and an overview of the available data per group and generation is shown in Table 1.

### Longevity is transmitted in the case group and not in the control group

Focusing on the original approach (Figure 1A), we determined to what extent longevity is transmitted in the original case and the control group by estimating SMRs per generation for all cases and controls separately. Table 2 shows that F1 cases had a similar survival pattern to birth cohort members of the same sex, indicating that they resemble a representative group of random Dutch persons aged ≥ 80 years and born between 1860 and 1875. The SMR for the descendants of the cases (F2 case descendants) was 0.87 (95%CI=0.84-0.89), indicating 13% less deaths than expected based on individuals from a similar birth cohort and sex. From here we refer to this as 13% excess survival (or, if appropriate, excess mortality) compared to the general population. The descendants of controls (F2 control descendants) had a similar survival pattern to the general population (SMR=1.01 (95%CI=0.96-1.05)). The spouses of the F2 case and control descendants surprisingly also showed a pattern of excess survival (SMR_caSe_F2spouses_=0.89 (95%CI=0.85-0.94) and **SMR_control _F2spouses_** = 0.9 (95%CI=0.83-0.97)). Next we observed 14% (95%CI=11%-16%) excess survival compared to the general population for F3 descendants of the F1 cases, whereas F3 control descendants resembled the general population (SMR=0.96 (95%CI=0.93-1.00)) just as observed in the F2 generation. The spouses of both F3 groups resembled the general population **(SMR_case_F3spouses_=1.00 (95%CI=0.95-1.05) & SMR_COntroLF3spouses_=1.07 (95%CI=0.99-** 1.15)). We conclude that two descendant generations of cases, who belong on average to the top 10% survivors, have 13-14% excess survival compared to the general populations and that the descendants of controls resemble the general population.

**Table 2:** Standardized mortality ratios for original case and control group individuals

| Role | Case group |  | Control group |  | Adjustment for right truncation |
| --- | --- | --- | --- | --- | --- |
|  | SMRs | Number (N) | SMRs | Number (N) |  |
| F1 IPs | 1.06 (0.99-1.13) | 884 | NA | NA | 80 years |
| F2 descendants | 0.87 (0.84-0.89) | 4416 | 1.01 (0.96-1.05) | 2203 | No adjustment |
| F2 spouses | 0.89 (0.85-0.94) | 1516 | 0.9 (0.83-0.97) | 697 | 20 years |
| F3 descendants | 0.86 (0.84-0.89) | 9015 | 0.96 (0.93-1.00) | 4353 | No adjustment |
| F3 spouses | 1.00 (0.95-1.05) | 2081 | 1.07 (0.99-1.15) | 1097 | 20 years |

Original cases (F1 IPs) died at 80 years or older, original controls (F1 IPs) died between 50 and 69 years. If persons could not die before a specific age due to direct or indirect selection, due to for example that all persons in a group were selected to have a child an adjustment for right truncation was applied so that a fair comparison could be made with their birth cohort members. An SMR for F1 control IPs could not be estimated due to a combination of left and right truncation in the data. The lifetables can only be adjusted for right or left truncation, but not a combination between the two.

To explore to what extent the survival of F2 and F3 descendants depends on the extremity of the longevity of their parents, we calculated SMRs for F2 and F3 case and control descendants with increasing parental longevity (for example, a parent belonged to the top 10%, 5%, or 1% survivors). We observed that the SMR decreased in descendants when defining parental longevity in terms of more extreme survival percentiles. This was the case for descendants of both the IP cases and controls although the effects were stronger in the descendants of the cases, especially in F3, since this group is now selected to have long-lived parents and grandparents (Supplementary table 1). This illustrates that selection on single long-lived persons belonging on average to the top 10% survivors, as we did for the IP selection, leads only to a modest transmission of longevity in two generations (max 14%). Likely, the control group includes misclassified persons of which the descendants do live longer, whereas the case group includes long-lived persons that do not transmit longevity to their descendants (potentially these are phenocopies). Such misclassification can jeopardize genetic studies immensely. To be able to evaluate living persons as potential carriers of the heritable longevity trait in genetic studies, we constructed and validated a familial longevity score.

### Constructing the Longevity Relatives Count score

We now look at the HSN data from a different perspective, the combined approach (Figure IB). In the combined approach we consider the F3 generation as the focal point of the pedigree, instead of the F1 generation, as was the case in the original approach. To identify individuals with the heritable longevity trait, we constructed the LRC score.

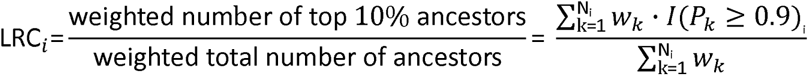

Where k=1,…,N_i_ are all the available ancestral blood relatives (from here: ancestors) of F3 descendant i used to build the score (parents, aunts and uncles and grandparent of the F3 descendants, Figure 1B), P_k_ is the sex and birth year-specific survival percentile, based on lifetables, of ancestor k, and l(P_k_ > 0.9) indicates if ancestor k belongs to the top 10% survivors. 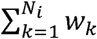 is the weighted total number of ancestors of F3 descendant i. The relationship coefficients are used as weights w_k_. The LRC score indicates the proportion of ancestors that has become long-lived. For example, an LRC of 0.5 indicates 50% long-lived ancestors (see methods for a more detailed and general description of the LRC score).

### Longevity is transmitted when at least 20% of all ancestors are long-lived

To determine what proportion of long-lived ancestors could be associated with the survival of F3 descendants, we calculated LRC scores for all F3 descendants and subsequently defined 9 mutually exclusive LRC groups (g) of F3 descendants: LRC_g1=0, LRC_g2=[≥0 & <0.1], LRC_g3=[≥0.1 & <0.2], LRC_g4=[≥0.2 & <0.3], LRC_g5=[≥0.3 & <0.4], LRC_g6=[≥0.4 & <0.5], LRC_g7=[≥0.5 & <0.6], LRC_g8=[≥0.6 & <0.7], LRC_g9=[≥0.7 & ≥1.0]. For each group of F3 descendants we explored whether they have a survival benefit compared to the general population by estimating SMRs (Figure 2). F3 descendants without any long-lived ancestors (LRC score of 0) had a survival pattern that resembled the general population (SMR=0.97 (95%CI=0.93-1.01)). Similarly, we observed a survival pattern that resembled the general population for F3 descendants with up to 20% long-lived ancestors (group 2 and 3, SMR=0.97 (95%CI=0.91-1.04) and SMR=0.95 (95%CI=0.91-1.00) respectively). This shows that the long-lived ancestors of group 2 and 3 F3 descendants were likely phenocopies instead of genetically enriched long-lived persons. We observed a pattern of excess survival for F3 descendants with more than 20% long-lived ancestors. The weakest significant effect was observed for group 3, with an SMR of 0.84 (95%CI=0.80-0.89) which is comparable to the excess survival of the F3 descendants of the singleton F1 cases in the original approach (first part of the results). The strongest significant effect was observed for group 8, with an SMR of 0.56 (95%CI=0.45-0.69). Hence, the higher the degree of long-lived ancestors, the lower the SMR. This indicates that the more long-lived ancestors an F3 descendant has, the higher the level of excess survival of these F3 descendants is compared to the general population, and the more likely that genetic effects drive the transmission of longevity.

**Figure 2:**
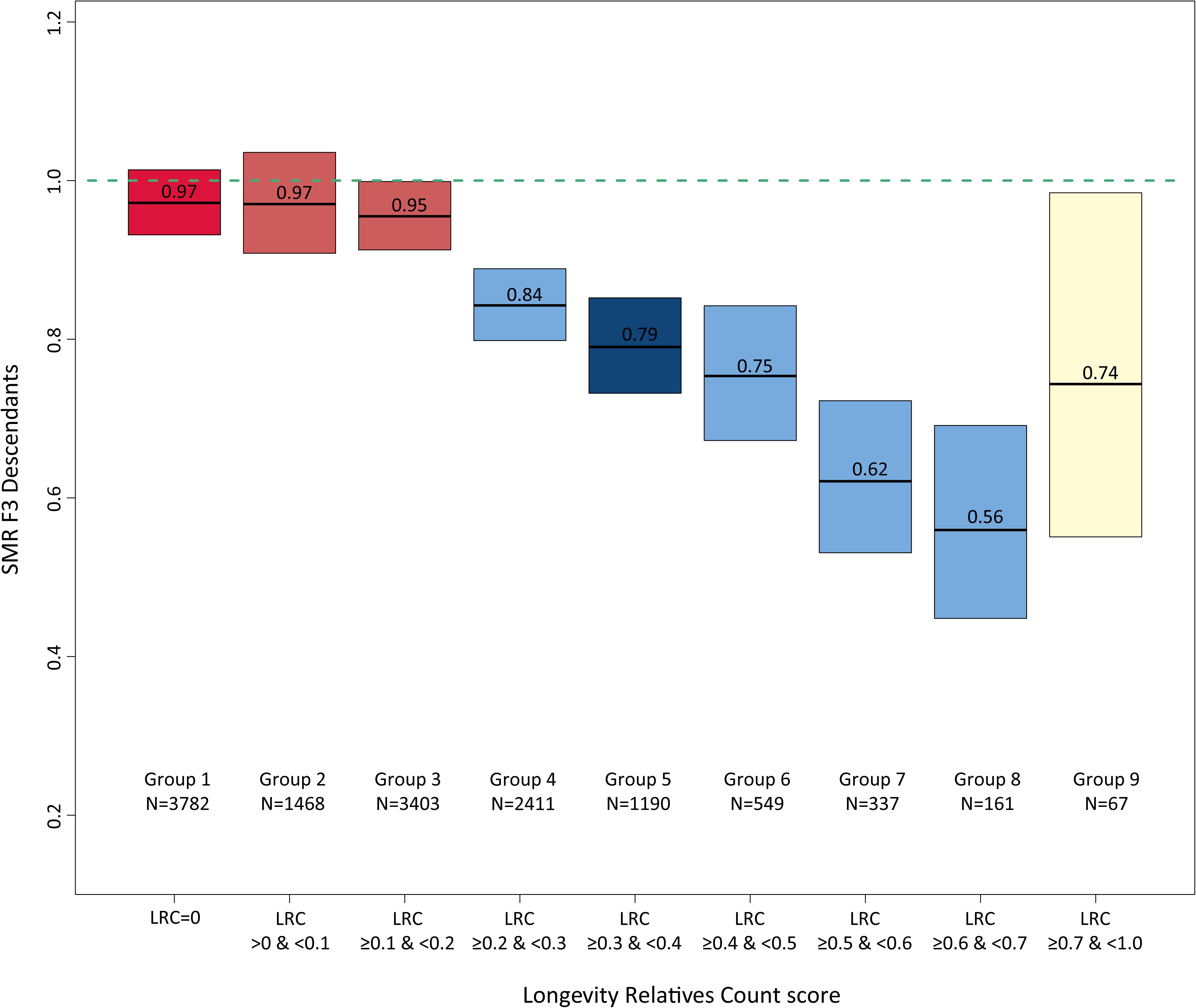
LRC score in mutually exclusive F3 descendant groups. The figure shows Standardized Mortality Ratios for all F3 descendants without missing mortality information. The F3 descendants are grouped into mutually exclusive groups based on the Longevity Relatives Count (LRC) score. The LRC score represents the family approach as illustrated in figure 1B. The dark red color of group one represents F3 descendants without any long-lived (top 10%) ancestors and are denoted as family controls. The light red represents F3 descendants who had more than 0 and less than 20% long-lived ancestors. The light blue colors represent the F3 descendants with 20% or more long-lived ancestors. The dark blue color represent our cut-off point for the family case definition. Hence all F3 descendants with 30% or more long-lived ancestors were considered family cases. The beige color of group 9 shows that this bar represents all F3 ancestors with more than 70% long-lived ancestors as their sample size was very low, we grouped them into one group.

Using the LRC score family method we defined a new case and control group in the F3 generation, which is based on the presence or absence of longevity among the ancestors of the F3 generation and potential excess survival or mortality in the F3 generation itself (Figure 1B). The F3 family controls include all F3 descendants without any long-lived ancestors (LRC score of 0, N=4,166). To define the F3 family cases we chose an LRC cutoff based on a trade-off between the size and the uncertainty, given by the sample size, of the SMR. The F3 family cases include all F3 descendants with at least 30% long-lived ancestors (LRC score ≥ 0.30 (N=2,526)). Even if F3 family cases are not long-lived themselves, their survival reflects the presence of longevity of their ancestors, which is transmitted by their parents. Similarly, F3 controls reflect the absence of longevity of their ancestors. Supplementary Figure 1 shows the variation in lifespan of the F3 family case and control descendants. F3 descendants with more than 0% and up to 20% long-lived ancestors (LRC score >0 and < 0.2) did not express excess survival (N=5,340). The F3 descendants with an LRC score ≥0.2 and < 0.30 showed some excess survival compared to the general population, but the size of the SMR was considered too low to enter our family case definition. Hence, we denoted them as non-classified (N=2,639).

### Strong survival advantage and genetic enrichment for F3 family cases

To validate the LRC score, we investigate survival differences, measured as age at death or last observation, between the F3 family cases and controls and used a Cox-type random effects (frailty) regression model to adjust for within-family relations of the F3 descendants. Figure 4 and table 3A show that F3 cases have a 25% (95%CI=18-31%) lower hazard of dying than F3 controls, even after adjustment for sibship size, birth year, and sex. The difference between the cases and controls became increasingly more pronounced when confining the cases to a higher proportion of long-lived ancestors, for example an LRC score of 0.40, 0.50, or 0.60, reflecting 40%, 50%, or 60% long-lived ancestors (Supplementary figure 2). The strongest effect was observed for those with an LRC score ≥ 0.60 (hazard ratio (HR) of 0.62 (95%CI=0.50-0.77)). The mortality pattern for the spouses of these F3 cases resembled that of the F3 controls (HR=0.94 (95%CI=0.82-1.07),Table 3B) and the general population (SMR=0.92 (95%CI=0.83-1.02)). The survival of the spouses, equal to the F3 controls and the general population, in addition to the absence of effects of environmental covariate adjustment, indicates that environmental factors were likely of limited influence to the observed survival benefit of the F3 cases as defined by our novel family based definition. Hence, the observed survival benefit of F3 cases likely represents a genetic longevity component.

**Figure 4.**
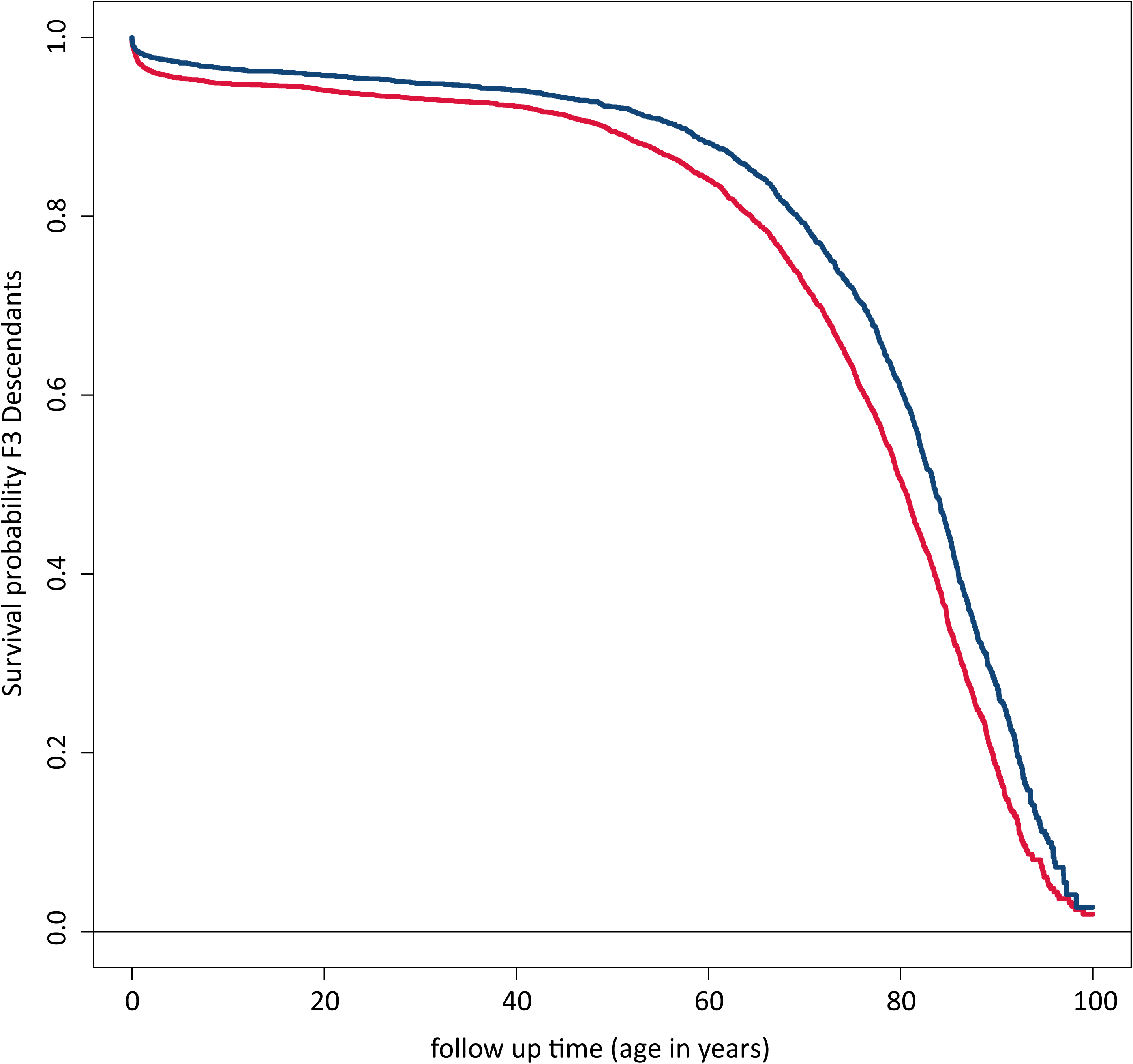
: Survival differences between family based cases and their spouses. This figure shows the survival curve for the difference in survival between the F3 family cases and controls. The figure is connected to Table 3A which shows the Hazard Ratios corresponding to the difference between the two curves. Blue color represent the cases, red color represents the controls.

**Table 3:** Mortality difference between family cases and controls and their spouses

|  | **A** |  |  | **B** |  |  |
| --- | --- | --- | --- | --- | --- | --- |
|  | **N (mean)** | **HR (95% CI)** | **P-value** | **N (mean)** | **HR (95% CI)** | **P-value** |
| Family based case/control group |  |  |  |  |  |  |
| Control group (ref) | 3714 (0.62) |  |  | 3714 (0.50) |  |  |
| Case group | 2282 (0.38) | 0.75 (0.69-0.82) | 1.75e-10 | 2282 (0.30) | 0.74 (0.68-0.80) | 4.08e-12 |
| Spouses of cases |  |  |  | 541 (0.07) | 0.94 (0.82-1.07) | 3.44e-01 |
| Spouses of controls |  |  |  | 937 (0.13) | 1.12 (1.00-1.25) | 4.07e-02 |
| Birth year | 5996 (1933) | 0.99 (0.98-0.99) | 1.99e-05 | 7474 (1932) | 0.98 (0.98-0.99) | 1.39e-12 |
| Sex |  |  |  |  |  |  |
| Males (ref) | 3133 (0.52) |  |  | 3364 (0.45) |  |  |
| Females | 2863 (0.48) | 0.56 (0.52-0.61) | <1.00e-15 | 4110 (0.55) | 0.49 (0.46-0.53) | <1.00e-15 |
| Sibship size |  |  |  |  |  |  |
| Small - 1-2 sibs (ref) | 1531 (0.26) |  |  |  |  |  |
| Medium - 3-5 sibs | 1770 (0.30) | 1.17 (1.04-1.32) | 8.51e-03 |  |  |  |
| Large - 6-8 sibs | 927 (0.15) | 1.22 (1.04-1.43) | 1.21e-02 |  |  |  |
| Exceptional - 9-15 sibs | 441 (0.07) | 1.36 (1.09-1.68) | 5.84e-03 |  |  |  |
| Single child - 0 sibs | 1327 (0.22) | 1.81 (1.62-2.02) | <1.00e-15 |  |  |  |

### Family cases live longer than those with one long-lived parent or grandparent

Next, we test if the F3 descendants with 30% long-lived ancestors (the family cases) have a stronger survival advantage than F3 descendants with at least 1 long-lived (top 10%) parent or grandparent. We actively selected this group of F3 descendants to have 1 long-lived parent or grandparent, meaning that other ancestors could also be long-lived but there was no active selection on the presence of their longevity (Supplementary Figure 3A and 3B), hence the designation ‘at least’ for this group. Subsequently, we tested if F3 descendants without long-lived ancestors (the family controls) had a similar survival pattern to the F3 descendants with parents resembling the general population (those with a grandparent who died between 40 and 59 years). Table 4 shows that we observed 14% (95%CI=11%-17%) excess survival compared to the general population for F3 descendants with at least one long-lived grandparent (F1). When identifying F3 descendants with at least one long-lived parent (F2), we observed 16% (95%CI=8%-24%) excess survival compared to the general population. Using the family method at 30% long-lived family members to identify F3 family cases, we observed 26% (95%CI=22%-30%) excess survival compared to the general population and this increased to 38% (95%CI=31%-45%) when applying a 50% threshold to the family method. For the identification of controls both methods seem to preform equally well, with almost identical SMRs of around 1. This indicates that the F3 controls, whether defined by having no long-lived ancestors or by grandparents dying between 40 and 50 years, have a similar survival pattern to the general population. We conclude that, at least for cases, the family method provides a better contrast in excess survival compared to the general population and seems to better represent the heritable longevity trait.

**Table 4:** Standardized Mortality Ratio for different F3 descendant groups

| Group | SMR | N |
| --- | --- | --- |
| Cases |  |  |
| F3 descendant with at least one long-lived grandparent | 0.86 (95%CI=0.83-0.89) | 4986 |
| F3 descendant with at least one long-lived parent | 0.84 (95%CI=0.76-0.92) | 852 |
| F3 descendant with $\geq 30\%$ long-lived ancestors (LRC $\geq 30\%$) | 0.74 (95%CI=0.70-0.78) | 2304 |
| F3 descendant with $\geq 50\%$ long-lived ancestors (LRC $\geq 50\%$) | 0.62 (95%CI=0.55-0.96) | 565 |
| Controls |  |  |
| F3 descendant with grandparent who died between 40 and 59 years | 0.96 (95%CI=0.93-1.00) | 4353 |
| F3 descendant with no long-lived ancestors (LRC = 0) | 0.97 (95%CI=0.93-1.01) | 3782 |

Long-lived is defined as belonging to the top 10% survivors of their birth cohort. Note that the group size (N) reflects only those with a known age at death as this was necessary to estimate a standardized mortality ratio.

Since the F3 descendants with ≥ 30% long-lived ancestors have a stronger survival advantage than those with at least one long-lived parent, it is possible to get an indication of how many F3 descendants did not appear to have a survival advantage compared to the general population, even though at least one parent was long-lived. This is relevant in view of case definitions used in large genetic studies into longevity. Figure 3 and Supplementary Figure 3 show that 919 F3 descendants had a long-lived parent. Out of those 919 F3 descendants, 247 (27%) had more than 0% but less than 20% long-lived ancestors (LRC > 0 and < 0.20) and thus as a group had an SMR that resembled the general population (Supplementary Figure 3D). The other 672 (73%) had exactly, or more than 20% long-lived ancestors (LRC ≥ 0.20) and thus, as a group, showed excess survival compared to the general population (Supplementary Figure 3B and C). These results suggest that if living persons are selected as case in genetic studies on the basis of one long-lived parent, 27% of these persons is unlikely to be a carrier of the longevity trait. Persons defined as 30% long-lived ancestors, on the other hand would be potential carriers.

**Figure 3:**
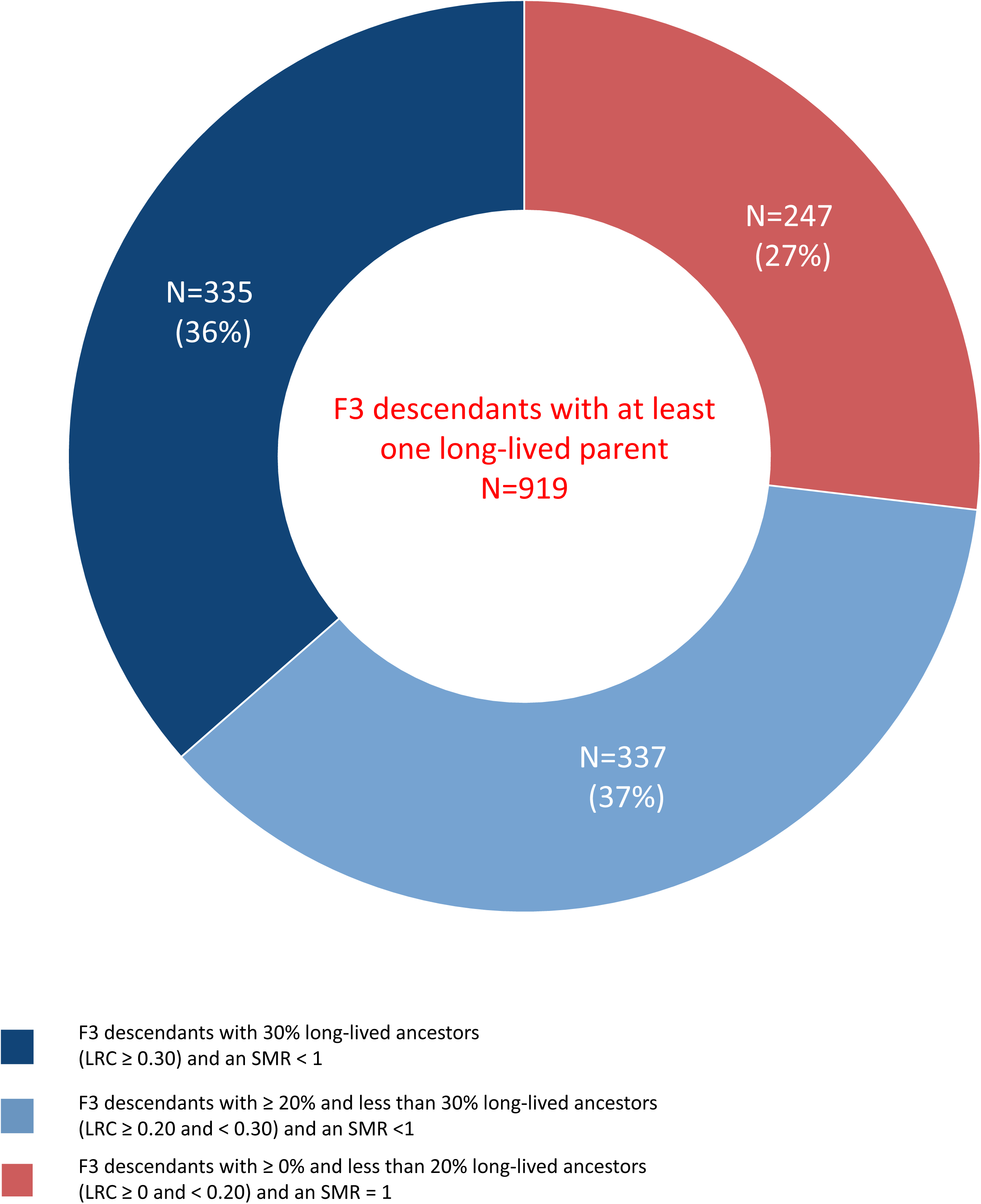
LRC score for F3 descendants with at least one long-lived parent. This center of this doughnut figure shows all F3 descendants (N=919) with at least one long-lived (top 10%) parent, ignoring the rest of the ancestors. Thus, at least means that they could have more than 1 long-lived ancestor but we actively selected for the presence of only 1 such ancestor. The edges of the doughnut illustrate the number and proportion of these 919 F3 descendants with at least one long-lived parent who had 1. 30% or more long-lived ancestors (LRC ≥ 0.30) and excess survival compared to the general population (SMR < 1), N=335 (36%) 2. between 20% and 30% long-lived ancestors (LRC ≥ 0.20 and < 0.30) and excess survival compared to the general population (SMR < 1), N=337 (37%) and 3. between 0% and 20% long-lived ancestors (LRC > 0.20 and < 0.20) and a similar survival pattern to the general population (SMR ∼ 1), N=247(27%).

## Discussion

Human longevity is heritable and clusters in specific families. Studying the familial clustering of longevity in these families is important to improve our understanding of genetic factors promoting longevity and healthy aging. The main observations supporting this are (1) In the original approach, we observed 14% excess survival of the cases compared to their birth cohort for two subsequent generations (F2-F3) while in the controls no such benefit was observed, (2) in the combined approach, the excess survival of the F3 cases compared to the general population was 26-38% depending on the proportion of long-lived family members being 30-50% and these estimates strongly overlap to the survival difference between the F3 family cases and controls based on the Cox models, (3) no excess survival as compared to the birth cohort and general population was observed for F3 controls, spouses of cases or controls and neither for F3 cases with up to 20% long-lived ancestors. The analyses in the HSN case/control dataset provides strong evidence that longevity is transmitted for at least 2 subsequent generations and only when at least 20% of all ancestors are long-lived. Moreover, the family cases seem to be genetically enriched for longevity while the controls resemble the general population. Finally, 27% of the F3 descendants showed a survival pattern similar to the general population even though they had at least one long-lived parent.

Previous family studies, usually focusing on 2 generations and single individuals, showed that siblings and children of long-lived persons lived longer than first degree ancestors of non-long-lived persons or population controls^5–7,9–15,45,49^. This knowledge about the familial clustering of longevity was utilized to construct longevity ranking scores such as the Family Mortality History Score (FMHS)^43^, the est(SE) which subsequently was developed into the FLOSS^39,44^, the Longevity Family Score (LFS) which is an adaptation to the est(SE) and the FMHS^15^, and finally a method was developed to rank individuals by the survival of their ancestors, the Familial Excess Longevity (FEL) score^38^. The FMHS, FLOSS, and LFS all resemble excess survival of a family (FMHS focus on parents and FLOSS and LFS focus on siblings) compared to the general population. The FEL score focuses on excess survival, defined as the difference between a person’s attained and expected age, derived from an accelerated failure time model. This excess survival was estimated for ancestors and from this a score was created for individuals. Although these scores all resemble a continuous familial estimate of a lifespan advantage and not necessarily longevity, they might be used as an inclusion tool for cases in genetic (association) studies^39^. However, these scores are not based on a clear longevity definition that represents the heritable longevity trait and they always require an arbitrary and difficult to interpret decision to make a cutoff in the scores so that they resemble longevity. In addition, the majority of the scores are not based on ancestors and thus do not capture the full family history of longevity. As such, the scores are not suitable to establish the proportion of family members that should be long-lived in order to properly define long-lived cases with a heritable longevity trait and thus, increase the power of genetic longevity studies.

To overcome these issues, we developed a novel tool based on mapping the longevity of a person’s ancestors, the LRC score. The LRC score can be used to select carriers of the heritable longevity trait (cases) and controls who resemble the general population. Another interesting group, which we did not address in this article, is composed of persons without any long-lived ancestors who themselves are long-lived. It may be interesting to study environmental factors contributing to a long and healthy life in this group. Here we used the LRC score to construct a novel family case and control group and observed a survival advantage for F3 case descendants, even when their parents were not necessarily long-lived, supporting the idea that a beneficial genetic component was transmitted. Likewise, the increase in the LRC score ≥ 20% associated with an increase in survival advantage for F3 descendants. This indicates that every additional ancestor contributes to the survival advantage of F3 descendants and confirms our previous findings in the LINKing System for historical demography (LINKS) data and the Utah Population Database (UPDB)^16^. This additive pattern is not readily expected if the observations are due to non-genetic factors, such as wealth, that cluster in families. The fact that none of the environmental confounders (sex, birth year, and sibship size) affected the survival differences between the family cases and controls provided additional evidence for the transmission of a genetic component. A final indication for the genetic enrichment of the family cases is based on the observed mortality pattern for the spouses of the family cases and controls which resembled the family controls themselves and the general population.

We observed that F3 descendants with at least one long-lived parent had less excess survival than a subset of these F3 descendants who had at least 30% long-lived ancestors and this difference increased when at least 50% of their ancestors were long-lived. These results indicate that some parents were long-lived but might not have transmitted their longevity to the subsequent F3 generation. In fact, 27% of the F3 descendants with at least one long-lived parent did not have an LRC ≥0.20 and, as a group, did not express excess survival. Hence the parents of theses 27% F3 descendants were sporadically long-lived as they did not transmit their longevity. Thus, genetic studies may benefit from a case definition, where cases are long-lived and have at least 30% long-lived ancestors, as current genetic studies, based on long-lived cases, often not include ancestral longevity in their case selection. Even though our data did not allow for an exact misclassification analysis, studies showed that the level of phenotypic misclassification in case and control annotation has a strong inhibiting effect on the power to identify variants in genetic association studies, including GWAS^42,50–58^. Moreover, it was shown that the power to identify genetic variants decreases at an equal rate to the level of misclassification^42^. For example, a study with 95% power to detect an association based on a sample of 100 cases and controls when there are no phenotypic errors may actually have only 75% power when 20% of the cases are misclassified as controls and vice versa^42^. Interestingly, when known, methods exist to adjust for the level of phenotypic misclassification^51–53,55,59^, providing opportunities for specific application in genetic longevity research.

Due to the nature of the HSN data we could not use the mortality data for the parents (F0), siblings (F1), and spouses (F1) of the F1 IPs. Mortality data was less incomplete for the F2 and F3 spouses (table 1A) but there was still a relatively large number of missing mortality data. Thus, for future studies with this dataset it might be interesting to extend the mortality information for these groups. Furthermore, life course data was only present for persons with an identified personal card or personal list (details in the methods section). Consequently, socio-economic status and religion was only available for a small part (around 15%) of the F3 descendants with an unequal share of availability between men and women. This led to the exclusion of these environmental factors from our analyses. Even though we could not adjust our models for socio-economic status and religion, it is known from other studies that those factors are not influencing the association between parental longevity and offspring survival^16^. Similarly, previous studies showed only a minor^60^ or no^16, 61^ influence of early and mid-life environmental covariates, such as farm ownership, parental literacy, parental and own occupation, and birth intervals, on the association between parental longevity and offspring survival. We, however, cannot completely rule out that other, unobserved non-genetic familial effects may affect our results. The observed excess survival of F2 case and control group spouses in the original approach seem to be an exception, as we observed a survival advantage for both groups. This is likely a form of ascertainment bias because mortality data for this group was difficult to obtain in the Dutch Personal Records Database, leading to an overrepresentation of high ages at death. These observations add to the mixed results about whether spouses married to a long-lived person have a survival advantage themselves^7, 11, 15–17, 62^.

Our results have two important implications. First, existing studies based on living study participants who have not yet reached the ages to express longevity, but have ancestral survival data, such as UK Biobank, can now better distinguish cases by incorporating a liability based on the LRC score. Second, new studies would obtain a maximum power to identify loci that promote survival to the highest ages in the population when cases are included with at least 30% (LRC≥0.30) ancestors who belong at least to the top 10% survivors of their birth cohort and are themselves among the 10% longest lived. More extreme selections can be made on the survival percentile by for example focusing on the top 5% or 1% survivors, and/or on the proportion of long-lived family members, for example 50%. However, this is not strictly necessary and might unnecessarily lead to limited sample sizes^16^. In addition, controls without any ancestors living to the top 10% survivors of their birth cohort should be included, as their mortality pattern resembles that of the general population. Finally, for future research it may be interesting to study the environmental factors causing the longevity in those individuals who were long-lived but had no long-lived ancestors. If our proposed method is consistently applied across studies, the comparative nature of longevity studies may improve and facilitate the discovery of novel genetic variants.

## Methods

### Historical Sample of the Netherlands

The Historical Sample of the Netherlands (HSN) Dataset Life Courses, Release 2010.01 is based on a sample of birth certificates and contains complete life course information for 37,137 Dutch individuals (index persons (IPs)) born in and between 1850 and 1922^46–48^. These 37,137 persons were subsequently identified in the Dutch population registers and followed in the registers throughout their entire life course^47, 48, 63^. The database includes information about the IPs’ household, including their siblings, parents, and children, occupation at several points in time and religion. Households were only followed as long as the IP was present in that household meaning that information on kin was only partly covered^48, 63^. For this study we selected 884 IPs who died at 80 years or beyond (case group) and 442 IPs who died between 40 and 59 years (control group), representing 1,326 disjoint families. IPs from both groups were born between 1860 and 1875. The case group was defined so that we would obtain a sample with overrepresentation of long-lived individuals. This was interesting since it would potentially allow to select on more extreme ages at death and still guarantee numbers reasonably large. The control group was selected to represent the mortality pattern of the general population of that time as best as possible. Individuals from both groups were selected to have an available date of birth, date of death, and at least one child should be identified. In conclusion, we identified 1,326 IPs (cases and controls), their F0 parents (N=2,652), F1 siblings (N=5,179), F2 descendants (N=7,404) and FI spouses (N=1,409), covering 3 filial generations (F0 - F2) spanning from 1788 to 1941 (Figure 1A and Table 1). The underlying data for this specific study were released as Kees Mandemakers and Cor Munnik, Historical Sample of the Netherlands. Project Genes, Germs and Resources. Dataset LongLives. Release 2016.01.

### Extending the HSN study

For this study we extended the pedigrees until we identified the living descendants for all 1326 families. From the population registers we know the names of all F2 descendants and we subsequently identified the F2 descendants on personal cards (PCs) and personal lists (PLs) which were obtained from the Dutch central bureau of genealogy (CBG). These PLs and PCs were respectively introduced in 1939 and 1994 as the individualized and subsequently, digitized form of the population register^48^. The cards contain similar information to the population registers and because of privacy legislation could only be obtained for deceased persons, one year after they passed away (https://cbg.nl/bronnen/cbg-verzamelingen/persoonskaarten-en-lijsten). Hence, from these cards we obtained similar life course and mortality information for the F2 descendants as for the F1 IPs and we obtained the names of their descendants (F3). We repeated this procedure until no cards could be obtained anymore, which was at the F3 generation. Thus the F4 generation was not identified on the PCs of PLs anymore. In conclusion, we identified and obtained information for the F2 descendants, F2 spouses, F3 descendants, F3 spouses, and F4 descendants (FigurelA and Tablel). We will refer to this database as the HSN case/control database.

### Obtaining information for the living descendants

In a final step we obtained as much mortality information as possible for the relatives of the identified persons and we obtained addresses, as contact information for the living descendants. This information was obtained through the Personal Records Database (PRD) which is managed by Dutch governmental service for identity information. https://www.government.nl/topics/personal-data/personal-records-database-brp. The PRD contains PL information on all Dutch citizens (alive and death) and PC information is continuously added. We were granted permission (permission number: 2016-0000364875) to obtain the date of death, date of last observation, current living address, and identifying information such as names of a person’s father and mother to double check if the person identified in the PRD was identical to the person in our HSN case/control database. Using the PRD we were able to obtain addresses for F3 and F4 descendants and additional mortality information for F2 descendants, F2 spouses, F3 descendants, F3 spouses, and F4 descendants (FigurelA and Tablel). The final database covers 57,337 persons from 1,326 five-generational families (F0-F4) and contains F1 index persons (IPs), their parents (F0), siblings (F1), spouses (F1), and 3 consecutive generations of descendants (F2-F4) and spouses (F2-F4), connecting deceased persons to their living descendants.

### Exclusion criteria and study population

Due to the nature of the source data there is a high rate of missing mortality information for FO parents, FI spouses and FI siblings, which we therefore excluded from analyses. We further excluded F4 descendants because 92% is still alive (Table 1 and Figure 1B). The final study population covers 37,825 persons from 1,326 three-generational families (F1-F3) and contains FI index persons (IPs), 2 consecutive generations of descendants (F2-F3) and 2 generations of spouses (F2-F3).

### Statistical analyses

Statistical analyses were conducted using R version 3.4.1^64^. We reported 95% confidence intervals (Cis) and considered p-values statistically significant at the 5% level (α = 0.05).

### Lifetables

In the Netherlands, population based cohort lifetables are available from 1850 until 2019^65,66^. These lifetables contain, for each birth year and sex, an estimate of the hazard of dying between ages x and x + n (hx) based on yearly intervals (n=1) up to 99 years of age.

Conditional cumulative hazards (Hx) and survival probabilities (Sx) can be derived using these hazards. In turn, we can determine to which sex and birth year based survival percentile each person of our study belonged to. For example: a person was born in 1876, was a female, and died at age 92. According to the lifetable information this person belonged to the top three percent survivors of her birth cohort, meaning that only three percent of the women born in 1876 reached a higher age. We used the lifetables to calculate the birth cohort and sex specific survival percentiles for all persons in the HSN case/control study. This approach prevents against the effects of secular mortality trends over the last centuries and enables comparisons across study populations^1, 14^. Supplementary Figure 6 shows the ages at death corresponding to the top 10, 5, and 1 percent survivors of their birth cohorts for the period 1850-1935.

### Standardized Mortality Ratios

To indicate excess mortality or excess survival of groups, such as F2 case or control group descendants in the HSN case/control study compared to Dutch birth cohort members of the same sex, we used Standardized Mortality Ratios (SMRs). An SMR is estimated by dividing the observed number of deaths by the expected number of deaths. The expected number of deaths are given by the sum of all individual cumulative hazards based on the birth cohort and sex specific lifetables of the Dutch population. An SMR between 1 and 0 indicates excess survival, an SMR of 1 indicates that the study population shows a similar survival to the reference population, and an SMR above 1 indicates excess mortality. The SMR can be estimated conditional on the specific age at which an individual starts to be observed in the study (correction for left truncation). This was necessary to avoid selection bias if individuals in a study population were not at risk of dying before a specific age of entry.

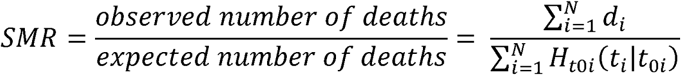

Where *d_i_*=dead status (1=dead, 0=alive), *H*_toi_=sex and birth year specific cumulative hazard based on lifetable, *t*[=timing, referring to age at death or last observation, *t_0i_*=liftable age conditioning, for example from birth (t*_0i_*=0), N= group sample size. Exact Cis were derived ^67^ and compared to bootstrap Cis for family data ^15^. Both methods provided identical Cis and thus, to reduce the amount of computational time necessary to estimate bootstrap Cis, we estimated exact Cis.

### Longevity Relatives Count score

Based on the results of a recent study which shows that longevity is heritable beyond the 10% survivors of their birth cohort and that multiple family members, such as parents and/or aunts and uncles, should belong to the top 10% survivors^16^ we constructed a novel score that summarizes the familial history of longevity, the Longevity Relatives Count score (LRC).

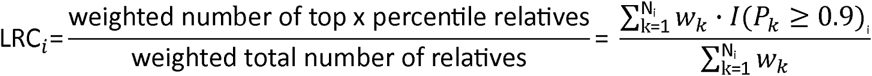

Where k=l,…,Ν_i_ are the available relatives of individual i used to build the score, P_k_ is the sex and birth year-specific survival percentile based on lifetables of relative k and l(P_k_ > 0.9) indicates if relative k belongs to the top 10% survivors 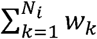 is the weighted total number of relatives of person i. The relationship coefficients are used as weights w_k_.. For example, persons share on average 50% of their nuclear DNA with their parents and siblings and this is 25% for aunts, uncles or grandparents. Hence, in the LRC, each parent and sibling contributes 0.5 to the score while each aunt, uncle or grandparent contributes only 0.25. This is consistent to a previous study of us, which shows that distant longevous relatives associate significantly, but less strong to a person’s survival than a close long-lived relative^16^. The higher the score, the higher the familial aggregation level of longevity. For example, a score of 0.5 indicates that 50% of a person’s relatives were long-lived. We utilized the LRC score to map the proportion of long-lived ancestors for all F3 descendants, select cases with the heritable longevity trait and controls resembling the general population, and compare the survival advantage of F3 descendants who had at least one long-lived parent to those who had at least 30% long-lived descendants. The LRC scores were based on all identified relatives of F3 descendants with sufficient data quality (Supplementary Figure 4 and 5).

### Survival analysis (Cox-type random effects regression model)

To investigate the extent of a survival difference between the family F3 case and control group we use a Cox-type random effects model:

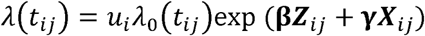

where t_ij_ is the age at death for person *j* in family *i*. λ_0_(t*_ij_*) refers to the baseline hazard, which is left unspecified in a Cox-type model, **β** is the vector of regression coefficients for the main effects of interest **(Z). γ** is a vector of regression coefficients for the effects of covariates and possible confounders **(X).** *U_i_ >* 0 refers to an unobserved random effect (frailty). In all Cox models we adjust for sibship size, birth year, and sex.

### Code availability

The scripts containing the code for data pre-processing and data analyses can be freely downloaded at: https://git.lumc.nl/molepi/PUBLIC/LRCscore.

### Data availability

Currently all data is cleaned and we are constructing a data description file. As soon as the data description file is completed the data will be made freely available in a data repository on DANS (https://dans.knaw.nl/en/front-page?setlanguage=en).

