## supplementary for "Longevity Relatives Count score identifies heritable longevity carriers and suggests case improvement in genetic studies"

Supplementary table 1: Increase in excess survival with an increase in parental survival percentile

| percentile threshold | SMR (CI) | Number (N) |
|---|---|---|
| **F2 children (original design - case)** | | |
| **No parental selection** | 0.87 (0.84-0.89) | 4416 |
| **≥85% surviving parent** | 0.85 (0.83-0.88) | 3770 |
| **≥90% surviving parent** | 0.85 (0.82-0.88) | 2565 |
| **≥95% surviving parent** | 0.86 (0.81-0.9) | 1336 |
| **≥99% surviving parent** | 0.80 (0.7-0.9) | 239 |
| **≥99.5% surviving parent** | 0.77 (0.65-0.9) | 130 |
| **F3 children (original design - case)** | | |
| **No parental selection** | 0.86 (0.84-0.89) | 9010 |
| **≥50% surviving parent** | 0.84 (0.82-0.87) | 7476 |
| **≥60% surviving parent** | 0.83 (0.81-0.86) | 6357 |
| **≥70% surviving parent** | 0.81 (0.78-0.84) | 5144 |
| **≥80% surviving parent** | 0.77 (0.74-0.81) | 3434 |
| **≥85% surviving parent** | 0.76 (0.72-0.8) | 2639 |
| **≥90% surviving parent** | 0.71 (0.67-0.76) | 1813 |
| **≥95% surviving parent** | 0.69 (0.63-0.76) | 798 |
| **≥99% surviving parent** | 0.65 (0.49-0.85) | 78 |
| **F3 children (original design - control)** | | |
| **No selection** | 0.96 (0.93-1) | 4353 |
| **≥50% surviving parent** | 0.95 (0.91-0.99) | 3425 |
| **≥60% surviving parent** | 0.92 (0.87-0.96) | 2828 |
| **≥70% surviving parent** | 0.89 (0.84-0.93) | 2168 |
| **≥80% surviving parent** | 0.85 (0.79-0.91) | 1479 |
| **≥85% surviving parent** | 0.84 (0.78-0.91) | 1107 |
| **≥90% surviving parent** | 0.84 (0.76-0.92) | 772 |
| **≥95% surviving parent** | 0.77 (0.67-0.89) | 317 |
| **≥99% surviving parent** | 0.94 (0.62-1.38) | 41 |

Surviving parent refers to the proband IP for the F2 children. It refers to having at least 1 parent belonging to the specified survival percentile threshold for the F3 children. Estimates are only based on the proband line and not on the spousal line.

Supplementary figure 1: Family cases and controls distributed by their reached age

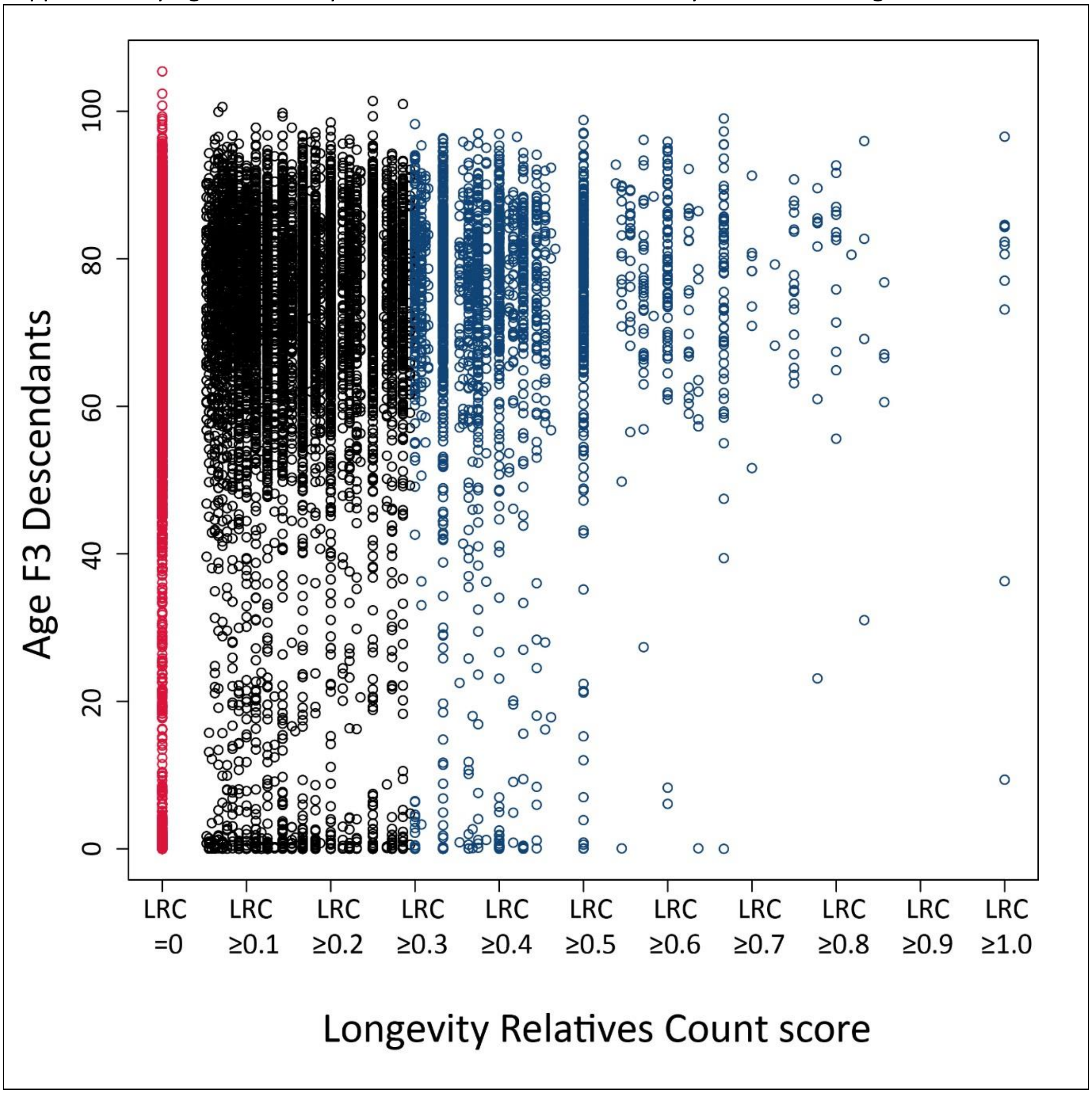


The x-axis represents the Longevity Relatives Count (LRC) scores of the F3 descendants. The y-axis represents the attained age of the F3 descendants. The attained age can either be an age at death or an age at last observation. Around 50% of the F3 descendants is still alive (table 1). The red color shows the family controls (those without any long-lived ancestors) and the blue color shows the family cases (those with an LRC ≥0.30).

Supplementary figure 2: SMR of F3 descendants with an increasing proportion of long-lived ancestors

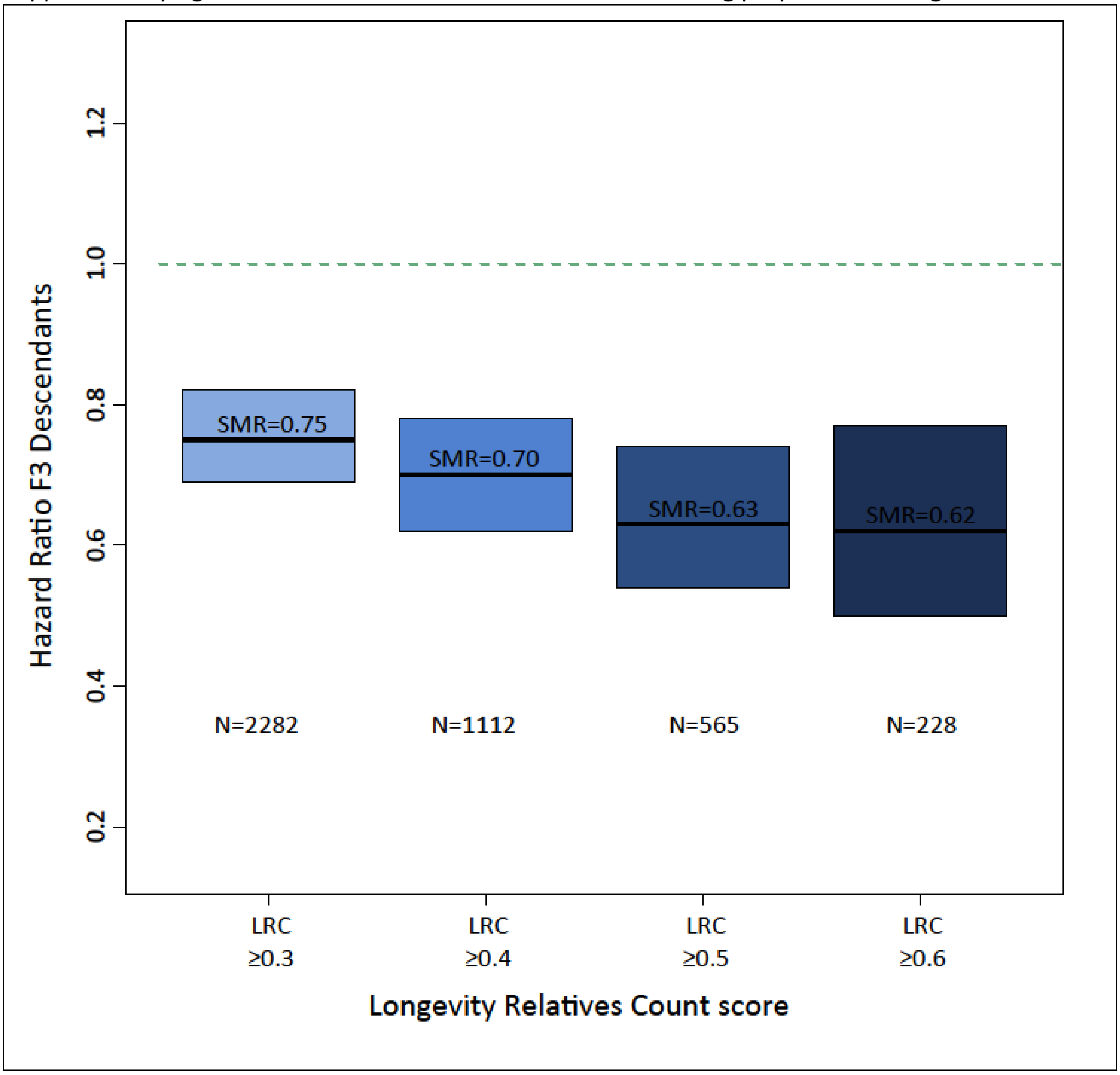


The x-axis represents the Longevity Relatives Count (LRC) score for the F3 descendants. The y-axis shows the Hazard Ratios (HRs) for the F3 descendants. The stronger the effect (lower HR) the darker the blue color.

Supplementary figure 3: Selection of F3 descendants based on the LRC and on at least one long-lived parent

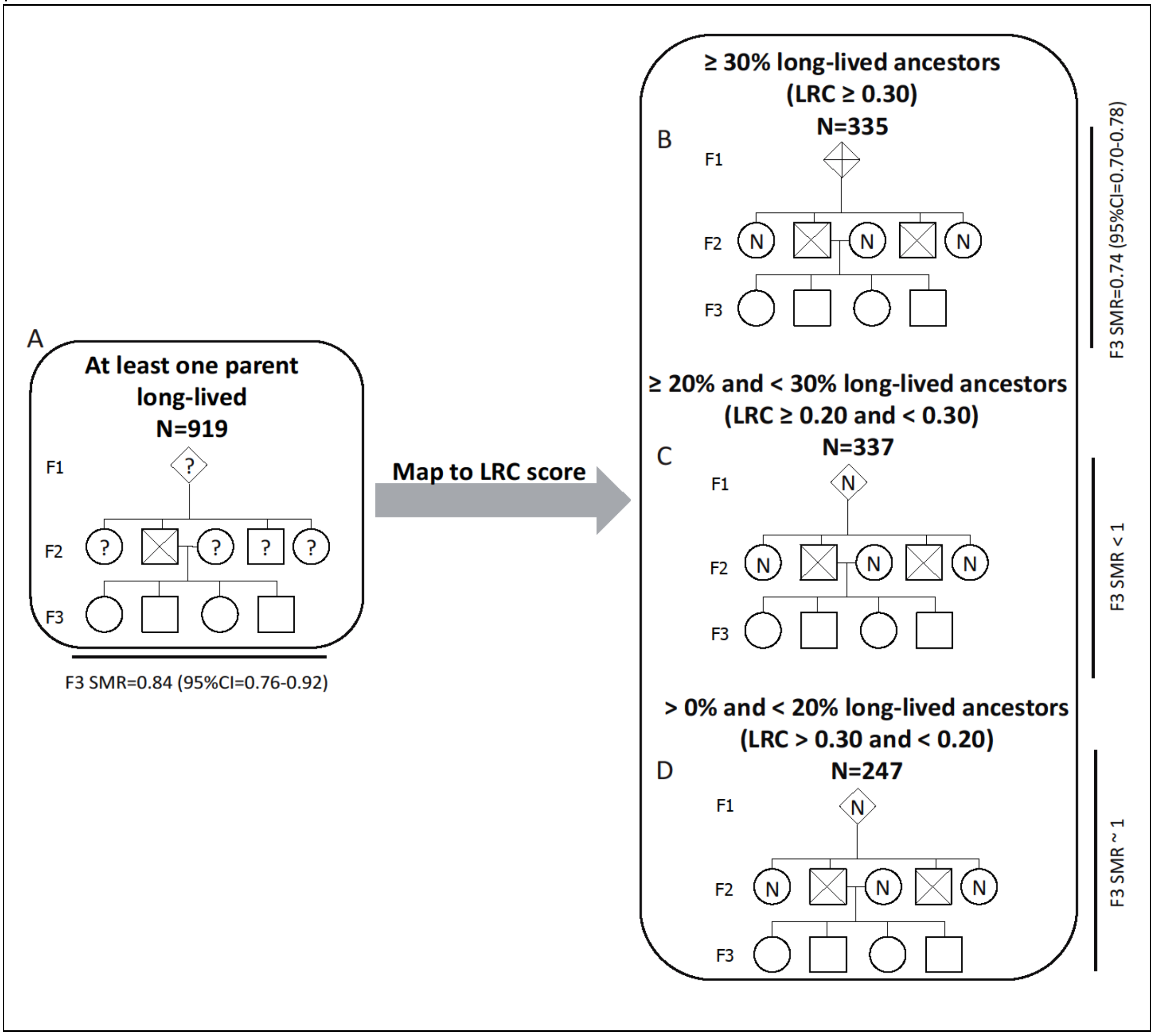


The ? sign shows that the survival of that specific ancestor was unknown. The N sign shows that the ancestor was nog long-lived (top 10% survivor). The X sign shows that the ancestor was long-lived. Panel A shows the F3 descendants with at least one long-lived parent. As illustrated, at least one means that we actively selected F3 descendants with one long-lived parent. That means that the other ancestors could also be long-lived but we did not take that information into account. This resembles the selection procedure of genetic longevity studies which focus on singletons. Panel B shows the ancestors with 30% long-lived ancestors or more and the corresponding standardized mortality ratio (SMR) observed for that group of F3 descendants. Panel C shows the F3 descendants who had between 20% and 30% long-lived ancestors and the corresponding SMR observed for that group. Panel D shows the F3 descendants with more than 0 and less than 20% long-lived ancestors and the corresponding SMR to that group.

Supplementary figure 4: Data cleaning procedure into the original and combined approach

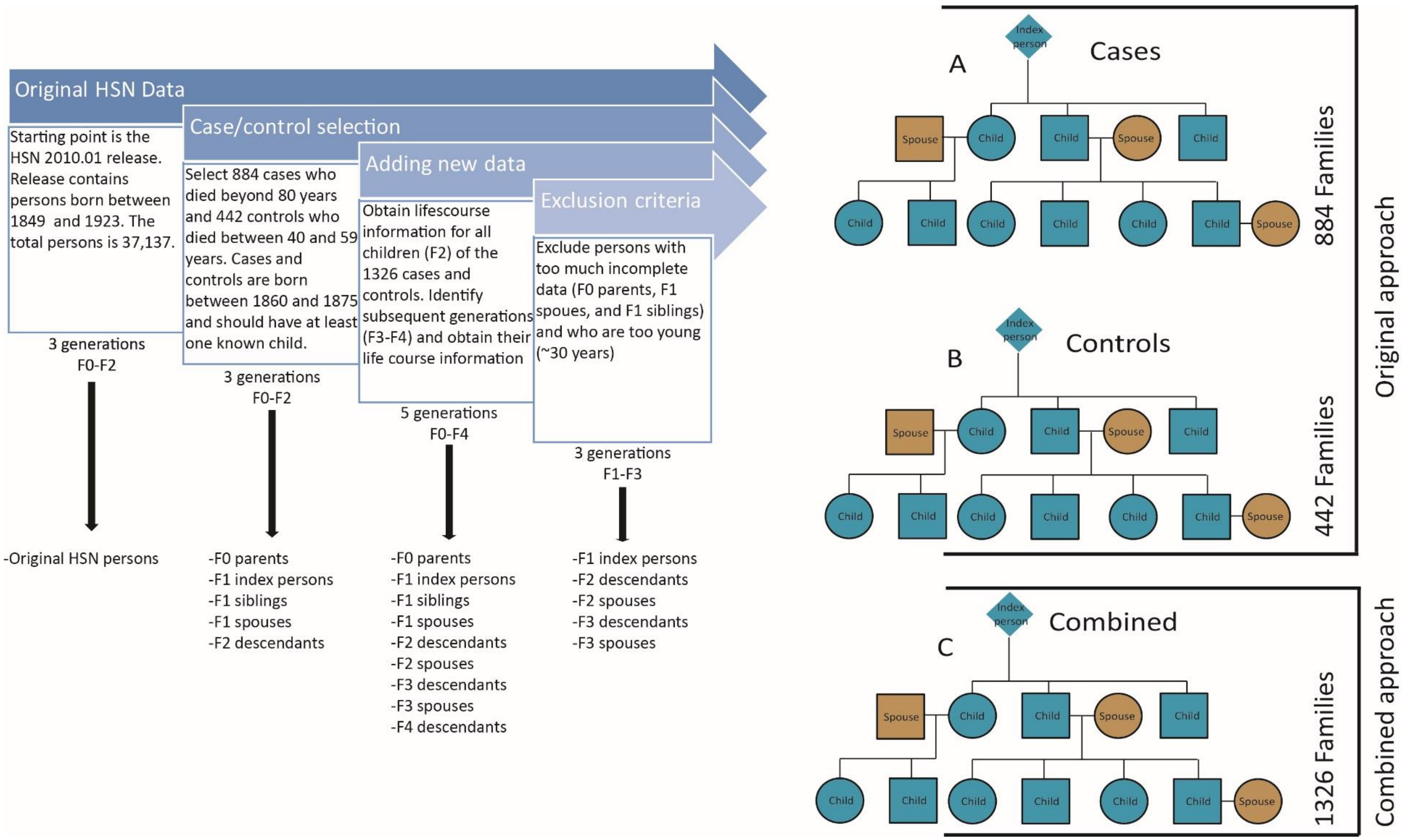


Figure provides an overview of the specific sample selection for this study. It starts with the original Historical Sample of the Netherlands data and works towards the specific study persons for this study and finally to the two different approaches used in this study.

Supplementary figure 5: Subsequent analyses steps in the original and combined approach

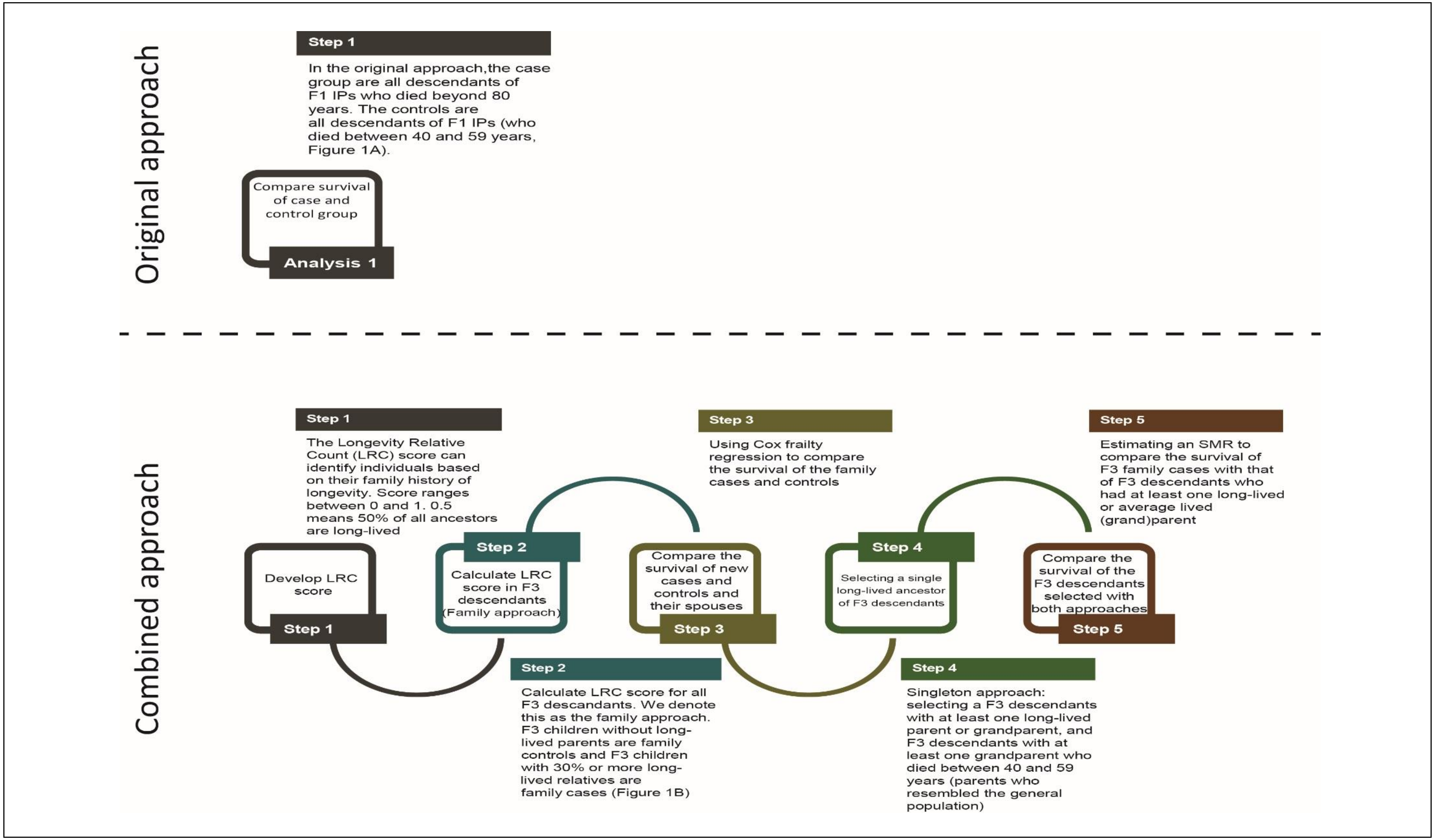


This figure shows the different steps used for the analyses in the original and the combined approach

Supplementary figure 6: HSN birth cohorts mapping of age by top 1, 5, and 10th percentile

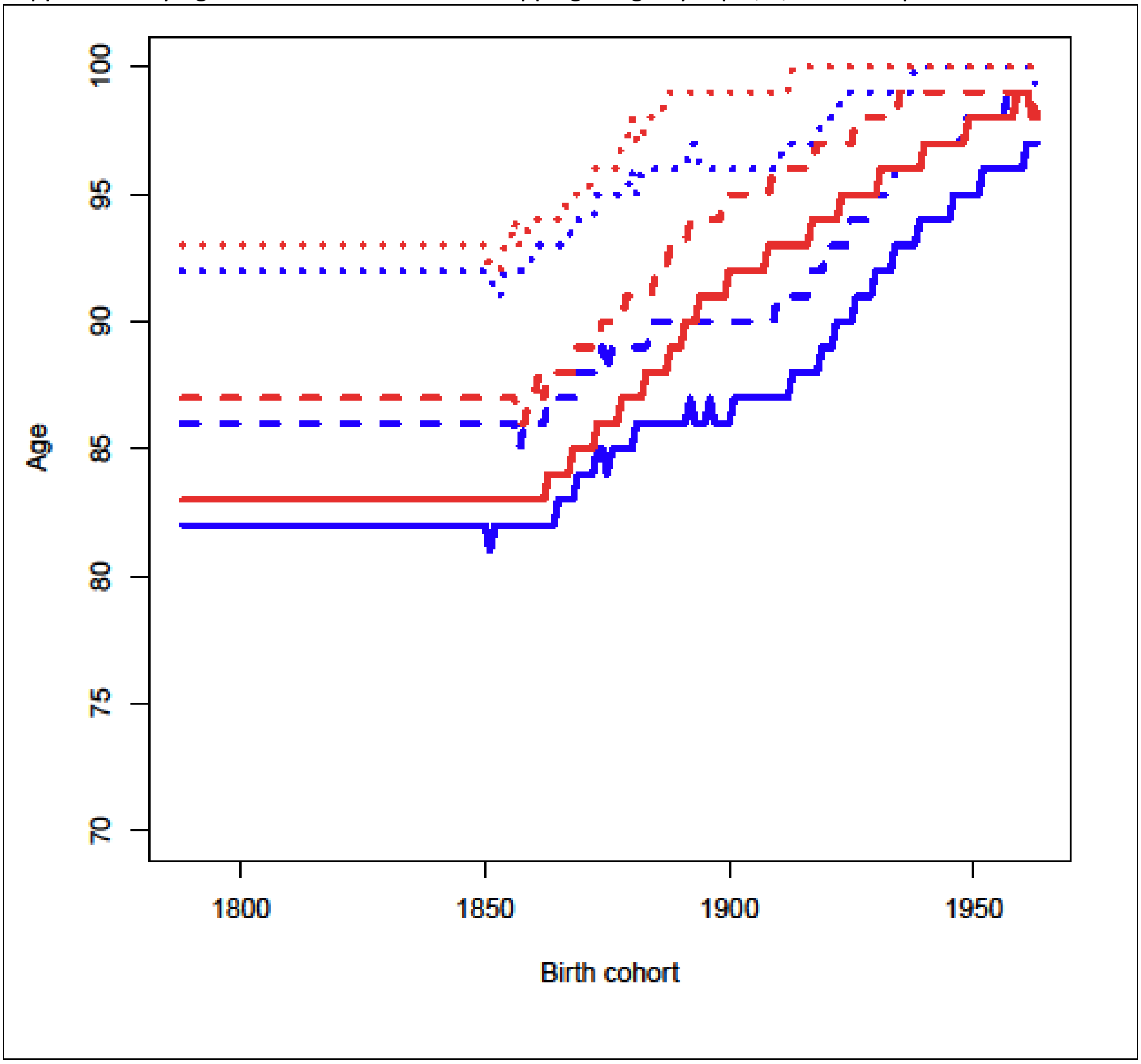


This figure represents the percentile-age pairings from the Dutch lifetables used to calculate survival percentiles in HSN case/control datasets. Line colors: Blue: men, Red: women. Line patterns: Dotted lines represent the top 1% survivors of the specific birth cohorts. Broken lines represent the top 5% survivors of the specific birth cohorts. Unbroken lines represent the top 10% survivors of the specific birth cohorts.
